# Transposable element insertions shape gene regulation and melanin production in a fungal pathogen

**DOI:** 10.1101/326124

**Authors:** Parvathy Krishnan, Lukas Meile, Clémence Plissonneau, Xin Ma, Fanny E. Hartmann, Daniel Croll, Bruce A. McDonald, Andrea Sánchez-Vallet

## Abstract

**Background**

Variation in gene expression contributes to phenotypic diversity within species and adaptation. However, very few cases of adaptive regulatory changes have been reported and the mechanisms underlying variation in gene expression remain largely unexplored. Fungal pathogen genomes are highly plastic and harbour numerous insertions of transposable elements, which can potentially contribute to gene expression regulation. In this work we elucidated how transposable elements contribute to variation of melanin accumulation, a quantitative adaptive trait of fungal pathogens that is involved in survival under stress conditions.

**Results**

We demonstrated that differential transcriptional regulation of the gene encoding the transcription factor Zmr1, which controls expression of the genes in the melanin biosynthetic gene cluster, is responsible for variation in melanin accumulation in the fungal plant pathogen *Zymoseptoria tritici*. We show that differences in melanin levels between two strains of *Z. tritici* are due to two levels of transcriptional regulation: 1) variation in the promoter sequence of *Zmr1*, and 2) an insertion of transposable elements upstream of the *Zmr1* promoter. Remarkably, independent insertions of transposable elements upstream of *Zmr1* occurred in 9% of *Z. tritici* strains from around the world and negatively regulated *Zmr1* expression, contributing to melanin accumulation variation.

**Conclusions**

Our studies demonstrate that different layers of transcriptional control fine-tune the synthesis of melanin. These regulatory mechanisms potentially evolved to balance the fitness costs associated with melanin production against its positive contribution to survival in stressful environments.

## Background

Understanding the genetic basis of adaptive traits is an important goal in ecology and evolutionary biology. Variation in gene expression is believed to underlie much of the phenotypic diversity within a species [1,2] and likely contributes to adaptive genetic variation [3]. However, most of the adaptive mutations identified until now are in coding sequences [4–6]. The reason for this is that protein variants are easier to identify because the genetic code enables *in silico* prediction of causative mutations.

Mutations outside of coding regions can lead to alterations in transcription, splicing, transcript stability or chromatin remodeling and consequently can affect the regulation of gene expression [6–8]. Modifications in cis-regulatory sequences, such as single nucleotide polymorphisms (SNPs) and indels, can affect their expression and are thought to be targets of adaptive evolution [9–14]. Additionally, insertions of transposable elements inside or outside the promoters may introduce elements that enhance or repress transcription and that induce changes in the chromatin state of adjacent regions, thus altering the expression of nearby genes [15–21]. Thus, transposable element insertions have the potential to contribute to phenotypic diversity through chromatin remodeling and regulation of gene expression.

In fungi, rapidly evolving regions, which frequently contain genes involved in virulence and stress tolerance, are often associated with transposable elements [22]. The contributions of transposable elements to evolution of adjacent regions and their effects on fungal diversity are frequently postulated [22–24], but have rarely been demonstrated. Many fungal plant pathogens are broadly distributed across the globe and are exposed to constantly fluctuating climatic conditions, a wide range of fungicides, and host immune defenses which can vary according to the host plant genotype [25,26]. Adaptation to changing environments typically favors the capacity to respond rapidly to stress. Additionally, populations that maintain a high standing variation for the genes that encode adaptive traits frequently are more successful in adapting to changes in the environment [27]. One such adaptive trait is melanization. Melanin is a broadly distributed secondary metabolite required by many fungi for host colonization and survival under stress conditions [28–31]. Two major types of melanin have been extensively described in fungi, namely dihydroxynaphthalene (DHN) and dihydroxyphenylalanine melanin [32–34]. A high diversity in melanin accumulation among individuals within a species provides a mechanism for differential adaptation and tolerance to rapidly changing and locally hazardous conditions [31].

*Zymoseptoria tritici* is a major wheat pathogen that has been extensively investigated for its adaptive potential to colonize different wheat cultivars and adapt to stressful conditions, including exposure to high temperatures and fungicides [35–37]. *Z. tritici* is known to have a plastic genome that includes numerous transposable element insertions (17% of the genome) and in which chromosomal rearrangements frequently occur [36,38,39]. It is thought that this genome plasticity can make important contributions to phenotypic variability, but the precise mechanisms underlying this phenomenon are not fully understood [36,37,40].

In four Swiss strains of *Z. tritici*, variable levels of melanin accumulation were observed and were postulated to contribute to differences in tolerance against abiotic stress, including fungicide resistance [33,41]. We aimed to further explore the genetic basis of differences in melanin accumulation by using a previously performed genetic mapping approach [33]. A single quantitative trait locus (QTL) was identified that contained part of the polyketide synthase 1 (*Pks1*) gene cluster, which is involved in biosynthesis of DHN melanin in other fungal species [32,34,42,43]. In this work we remapped the QTL to the genome of one of the parental strains and we obtained a narrower and shifted QTL confidence interval, which allowed us to determine the genetic basis of the differences in melanin accumulation. We show that variation in gene expression, instead of variation in the coding sequence, underlies the observed differences in melanin accumulation. Variation in expression of a single gene, encoding the homolog of the transcription factor Cmr1 (*Colletotrichum* melanin regulation 1), which we renamed Zmr1 (for *Zymoseptoria* melanin regulation 1), explained the variation in melanization. We discovered two independent causes of variation in gene expression, namely SNPs in the promoter of *Zmr1* and an insertion of transposable elements upstream of the *Zmr1* promoter. We then showed that diversity in melanin accumulation at the species level is determined in part by independent insertions of transposable elements, which regulate *Zmr1* expression. We hypothesize that the complex regulation of *Zmr1* facilitates the emergence and maintenance of diversity in melanization to optimize a trade-off between the deleterious effect of melanin on the growth rate and its favorable effects on survival in stressful environments.

## Results

### Differences in melanin accumulation are determined by the *Pks1* cluster

Melanin accumulation in the Swiss *Z. tritici* strain 3D1 was lower than in the strain 3D7 at early time points (10 days post inoculation, dpi). The differences in melanization were temporal, as the lighter strain 3D1 accumulated similar amounts of melanin as 3D7 at later developmental stages (11-12 dpi; Fig 1A, Additional file 1). We explored the genetic basis of these differences in melanin accumulation by using the previously obtained QTL for these two strains [33]. To narrow down the confidence interval, a new genetic map was obtained by using the completely assembled genome of the parental strain 3D7 [39] instead of the genome of the reference strain IPO323. This strategy provided us with approximately 10 times more SNP markers and enabled us to identify additional crossover events. The newly mapped 95% confidence interval of the melanization QTL was narrowed from 43429 bp to 18135 bp and contained six genes instead of twelve. The new QTL position shifted with respect to the earlier position, with an overlapping region of only 9299 bp. The region shared between the two QTLs contained the promoter of a gene encoding the homolog of the transcription factor Cmr1 *(Colletotrichum* melanin regulation 1), which we renamed Zmr1 (for *Zymoseptoria* melanin regulation 1). Two of the genes within the new confidence interval belonged to the *Pks1* cluster, namely *Zmr1* and 1,3,8-trihydroxynaphthalene reductase *(Thr1*, Fig 1B, Additional files 2 and 3).

**Fig. 1.**
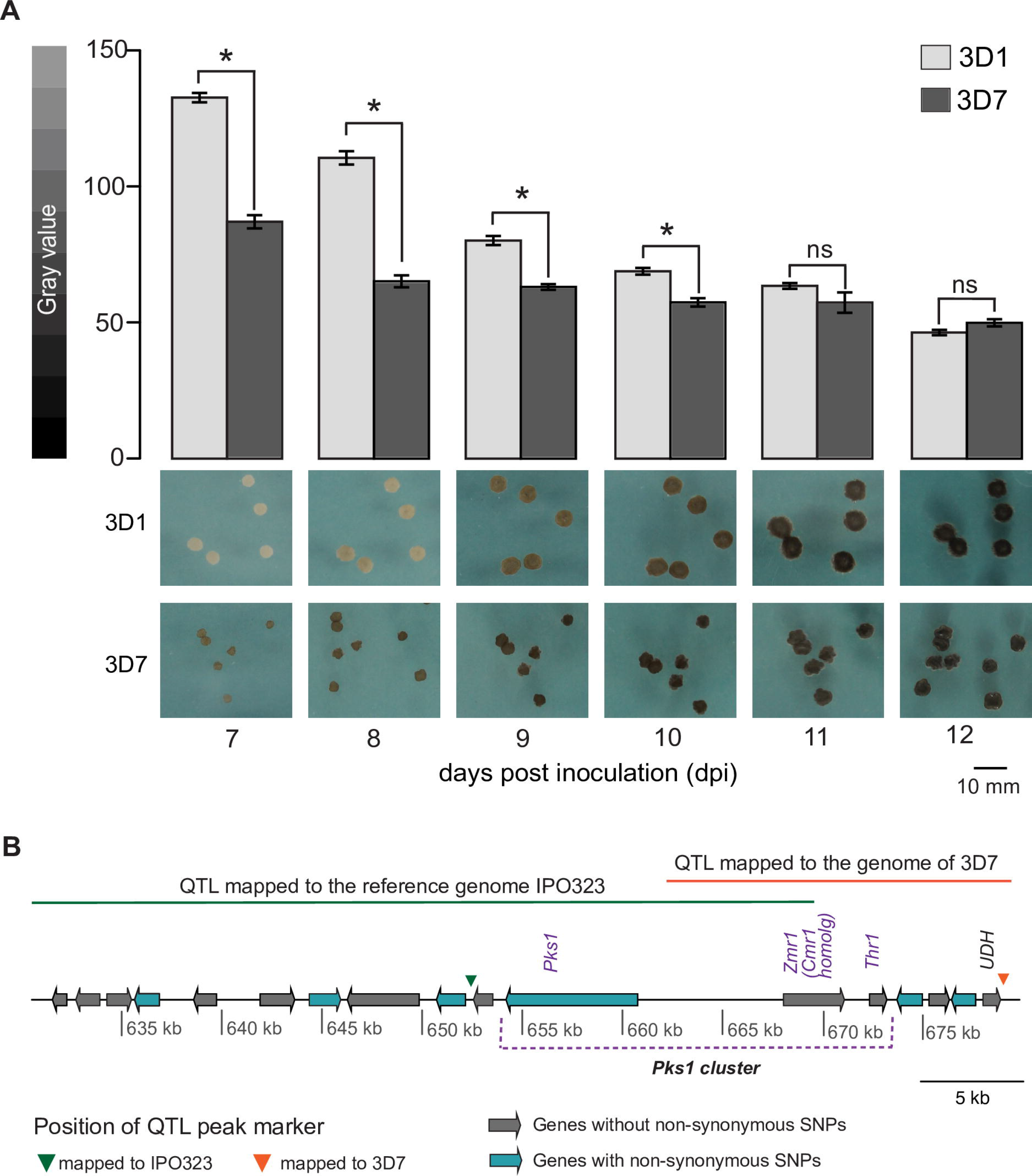
Variability in melanization levels between 3D1 and 3D7 maps to the *Pks1* cluster. (A) The Swiss strain 3D1 is less melanized than 3D7. Melanization levels of 3D1 and 3D7 at 7 to 12 days. Bars represent standard errors of the mean gray value based on at least 60 colonies. Asterisks indicate significant differences according to Kruskal-Wallis test (p-value ≤ 0.05; ns = non-significant). Representative pictures of both strains are shown below the bar plot for all the time points. The experiment was performed three times with similar results. Gray value scale (0 = black, 255 = white) is shown on the left. (B) Genes in the 95% confidence interval of the QTL mapped to the genomes of the reference strain IPO323 and of the parental Swiss strain 3D7. The shift in the position of the QTL, genes with and without non-synonymous mutations and the positions of the QTL peaking markers are indicated.

### A transposable element insertion in the *Pks1* gene cluster occurs only in the less melanized strain

The most obvious candidate genes to explain differential melanin accumulation in the two parental strains were *Zmr1* and *Thr1*. Both encoded proteins were identical between the parental strains (Additional file 3) and no mutations were detected in the promoter (1000 bp upstream of the start codon) of *Thr1.* However, twelve SNPs were identified in the promoter of *Zmr1* (Fig 2A) and we hypothesized that these SNPs could explain the differences in melanization. A comparison of the parental genomes revealed a loss of synteny in the QTL. We found an insertion of a transposable element island of approximately 30 kb, located 1862 bp upstream of the *Zmr1* start codon, only in the lighter strain 3D1 (Fig 2B). The sequences adjacent to the transposable element island, including the full *Pks1* gene cluster, showed a high conservation of synteny between the two parental genomes. The transposable element island consisted of 13 transposable elements and possessed both DNA transposons (of the TIR order) and retro-transposons (of the LTR and LINE orders) interspersed by simple repeats (Fig 2B).

**Fig 2.**
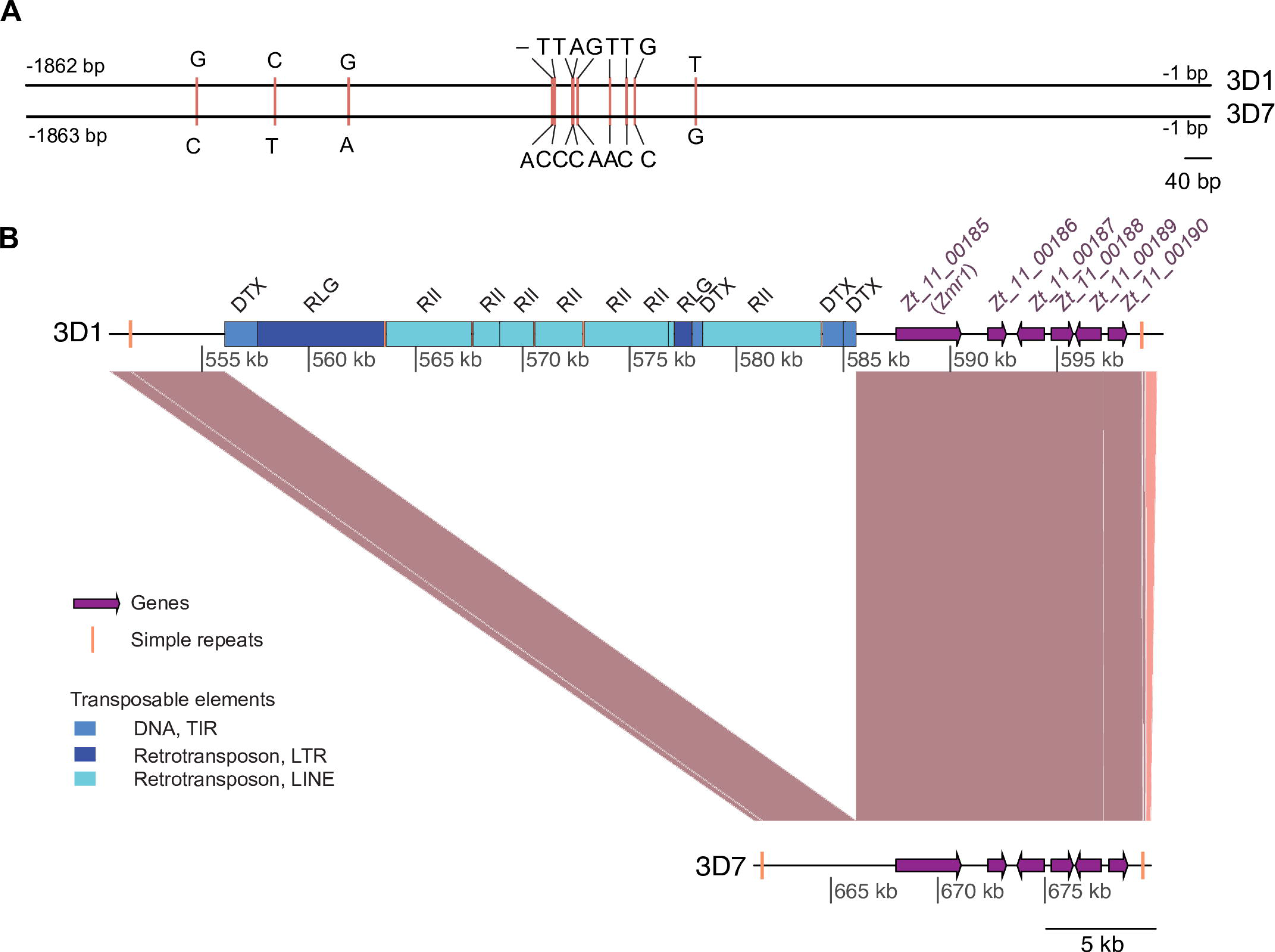
Differences between 3D1 and 3D7 in *Zmr1* regulatory sequences. (A) Alignment of the nucleotide sequences 1862 bp upstream of the coding sequence of *Zmr1* in 3D1 and 3D7. Horizontal pink bars indicate the 12 SNPs in the promoter region. (B) Synteny plot of the QTL between 3D1 and 3D7 showing the insertion of an island of transposable elements 1.8 kb upstream of the start codon of *Zmr1* in the lighter strain 3D1. Brown lines indicate collinear sequences. The positions of genes and transposable elements are shown using purple arrows and blue bars, respectively. Vertical yellow lines indicate simple repeats. The different shades of blue represent different classes of transposable elements that were classified according to the three-letter code described in Wicker *et al.*, (2016). The first letter indicates the class (R = RNA class and D = DNA class), the second letter indicates the order (L = LTR, I = Line, T= TIR) and the third letter indicates the superfamily (G = *Gypsy*, I = *I*, X = unknown).

### *Zmr1* expression is different between the two parental strains

We hypothesized that changes in non-coding regions could underlie natural variation in levels of melanization. Transposable element insertions upstream of the promoter and/or mutations in the promoter could lead to differential regulation of the genes in the *Pks1* gene cluster and, consequently, to different levels of melanin synthesis and accumulation. We found that *Zmr1* expression was higher in the darker strain 3D7 than in the lighter strain 3D1 at a time point when differences in melanin accumulation were detected (7 dpi). No significant differences in expression levels were observed at a later developmental stage (9 dpi), when the degree of melanization in 3D1 was higher (Fig 3, Additional file 4). Thus, we postulated that differential regulation of *Zmr1* expression, potentially mediated by differences in non-coding sequences, could underlie differences in this adaptive trait.

**Fig 3.**
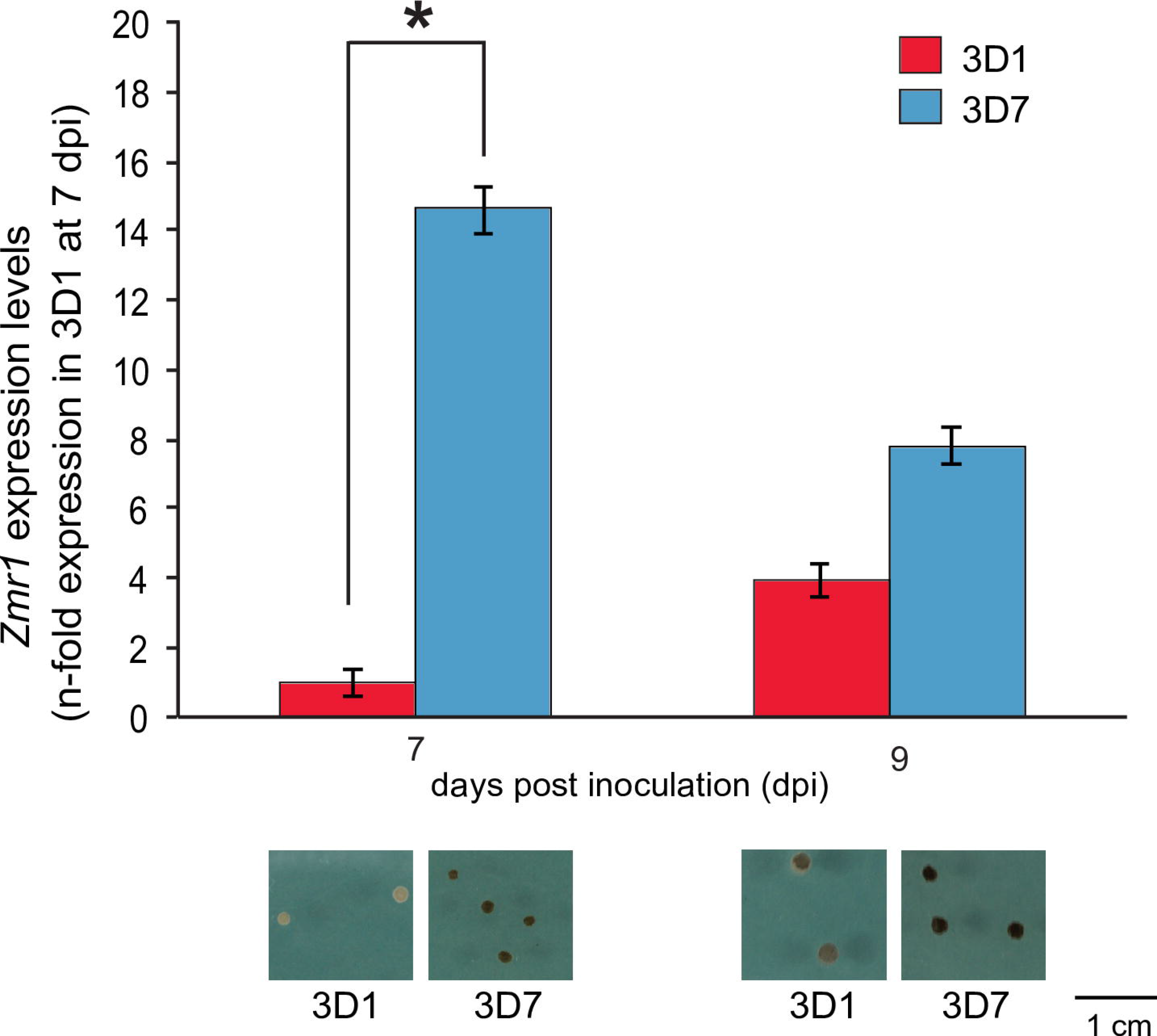
*Zmrl* expression is higher in the darker strain (3D7) compared to 3D1. *Zmr1* mean expression and standard deviation in 3D1 and 3D7 at 7 and 9 days post inoculation (dpi) relative to the expression of *Zmr1* in 3D1 at 7 dpi. Colonies grown on at least three different YMS plates were pooled for RNA extraction. The experiment was performed three times with similar results. Asterisks represent significant differences between 3D1 and 3D7 (p-value ≤ 0.05, Kruskal-Wallis test). A representative picture of each strain is shown at 7 and 9 dpi.

### Zmr1 regulates melanin biosynthesis in *Z. tritici*

To determine the role of Zmr1 in melanin accumulation in *Z. tritici*, we generated *Zmr1* knockout mutants by homologous recombination in strains 3D1 and 3D7 (Δ*zmr1*). No melanin accumulation was observed in Δ*zmrl* mutant colonies grown *in vitro* or in pycnidia formed on wheat leaves in both genetic backgrounds (Additional file 5), confirming that Zmr1 is required for melanin biosynthesis in *Z. tritici.* We further explored the function of the transcription factor Zmr1 in regulation of gene expression by pursuing a comparative transcriptomic analysis of the wild type strains and the Δ*zmr1* mutants, in both 3D1 and 3D7 backgrounds. Twelve genes were down-regulated in both Δ*zmrl* mutants (Table 1, Additional file 6). The expression levels of all the genes described to be involved in the DHN melanin biosynthetic pathway were significantly reduced. Remarkably, the expression of *Pks1* and *Thr1* was nearly abolished in the mutants (Table 1, Additional file 5 and 6). Transcriptomic profiling corroborated the hypothesis that Zmr1 is a major regulator of the genes involved in the DHN melanin biosynthetic pathway. We showed that DHN melanin is the only type of melanin accumulated in *in vitro* grown colonies and in *Z. tritici* pycnidia produced *in planta.*

**Table 1:** List of genes significantly down-regulated in *Zmr1* mutants in both 3D1 and 3D7 backgrounds.

| Gene ID | Log<sub>2</sub> FC |  | Annotation |
| --- | --- | --- | --- |
|  | 3D1 | 3D7 |  |
| **Zt09_11_00186** | **-9.92** | **-8.93** | **1,3,8-trihydroxynaphthalene reductase (*Thr1*)** |
| Zt09_4_00395 | -5.35 | -6.48 | Similar to cytochrome p450 |
| Zt09_2_00626 | -3.66 | -1.61 | Hypothetical protein |
| **Zt09_11_00184** | **-3.22** | **-4.35** | **Polyketide synthase (*Pks1*)** |
| Zt09_2_00074 | -2.5 | -5.09 | Predicted protein |
| **Zt09_3_00872** | **-2.2** | **-1.76** | **Homolog of *Aspergillus* yellowish green (*Ayg1*)** |
| Zt09_4_00393 | -2.19 | -3.27 | Hypothetical protein |
| Zt09_1_00117 | -2.15 | -4.14 | Hypothetical protein |
| Zt09_4_00394 | -2.15 | -2.4 | Salicylate hydroxylase |
| Zt09_3_00063 | -2.01 | -5.28 | Glycoside hydrolase family 18 protein |
| **Zt09_11_00185** | **-1.76** | **-1.94** | ***Zymoseptoria* melanin regulation 1 (*Zmr1*)** |
| Zt09_1_00074 | -1.76 | -3.82 | Hypothetical protein |
| **Zt09_1_00268** | **-1.67** | **-2.5** | Scytalone dehydratase (*Scd1*) |

Log_2_ fold change (log_2_ FC) expression values (counts per million mapped) of genes significantly down-regulated (Benjamin-Hochberg false discovery rates (FDR) ≤ 0.05 and adjusted p-value ≤ 0.05) in both 3D1Δ*zmr1* and 3D7Δ*zmrl*, compared to their respective wild type. In bold are genes already described to be involved in the DHN melanin pathway in other fungal species.

### Sequence variation in the promoter of *Zmr1* contributes to the differential regulation of *Zmr1*

We next postulated that the basis of the differential accumulation of melanin in 3D1 and 3D7 is the differential expression of *Zmr1* (Fig 3), which could potentially be caused by modifications in the promoter or by the transposable element insertion (Fig 2). The contribution of promoter modifications to *Zmr1* expression was evaluated by analyzing allele replacement lines in the 3D7 background. Increased melanization was achieved by *in locus* expression of both 3D1 and 3D7 *Zmr1* alleles in 3D7*Δzmr1* (3D7*Δzmrl* + *Zmr1*_*3D1*_, 3D7*Δzmr1* + *Zmr1*_*3D7*_) compared to the knockout, confirming the role of *Zmrl* in melanin biosynthesis in *Z. tritici.* Remarkably, although the 3D7 allele fully complemented the knockout phenotype, *in locus* expression of the 3D1 allele led to an intermediate phenotype between the knockout and the wild type (Fig 4, Additional file 7), suggesting that differential accumulation of melanin is caused by SNPs in the *Zmr1* promoter.

**Fig 4.**
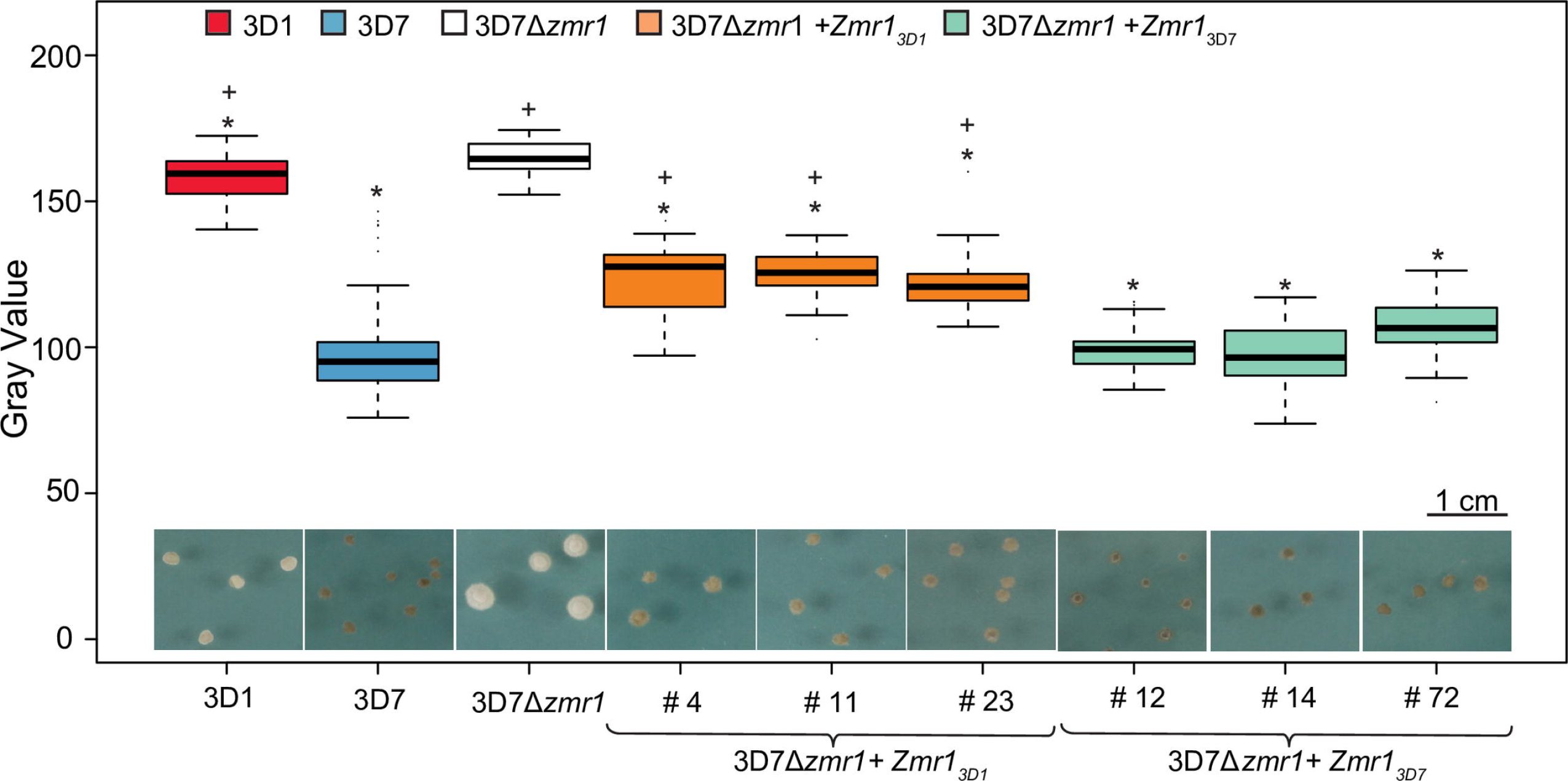
Sequence variation in the *Zmr1* promoter contributes to differences in melanin accumulation. Gray values for 3D1, 3D7, the *Zmr1* knockout in 3D7 (3D7*Δzmr1*), three *in locus* complementation transformants (3D7*Δzmrl* + Zmr1_3D7_ #4, #11, #23) and three *in locus* allele swap mutants of *Zmr1* (3D7*Δzmrl* + *Zmr1_3D1_* #12, #14, #72), all grown for 7 days. Asterisks (*) and plus (+) indicate significant differences in gray values of each strain with respect to the gray value of 3D7 *Δzmr1* and 3D7, respectively (p-value ≤ 0.05, Kruskal-Wallis). At least 20 colonies (replicates) grown on three different plates were evaluated. The experiment was performed twice with similar results.

### Insertion of a transposable element island upstream of the *Zmrl* promoter down-regulates *Zmr1* expression

We investigated whether the transposable element insertion in the lighter strain 3D1 modulated *Zmr1* expression. We made use of *Δzmr1* mutants of both 3D1 and 3D7, in which the *Zmr1* gene was disrupted by a hygromycin resistance cassette under the control of a constitutive promoter and of the ectopic controls, in which the hygromycin resistance cassette did not disrupt the *Zmr1* gene but was inserted elsewhere in the genome (Fig 5A, C). In the 3D7 background, the knockouts and the ectopic lines displayed similar growth in the presence of hygromycin (Fig 5B). Remarkably, we observed that growth of all three independent *Δzmr1* knockouts in the 3D1 background was lower than growth of the ectopic transformants in hygromycin-containing medium (Fig 5D). Hence, we hypothesized that the transposable element cluster silenced the expression of the hygromycin resistance gene, with the observed phenotype in the mutant likely reflecting the contribution of the transposable element insertion to *Zmr1* expression regulation.

**Fig 5.**
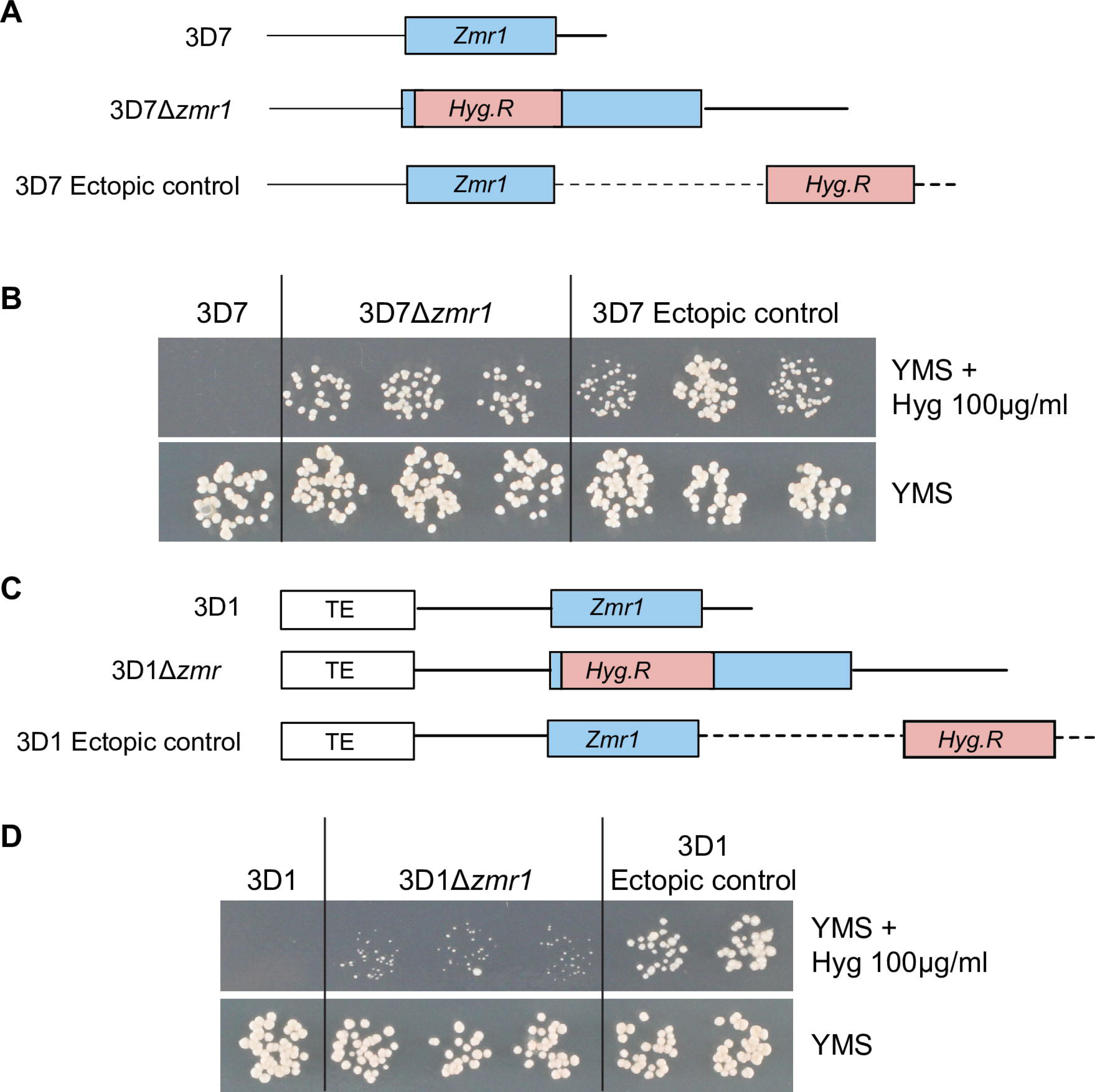
The transposable element insertion upstream of *Zmr1* influences the expression of adjacent genes. (A) Schematic representation of the *Zmr1* locus in the wild type 3D7 and the insertion of a hygromycin resistance cassette (*Hyg.R*) in 3D7*Δzmr1* and in the ectopic controls. (B) Growth of 3D7, 3D7*Δzmr1* and the ectopic controls in yeast malt sucrose (YMS) plates with and without hygromycin (100 μg/ml). (C) Schematic representation of the *Zmr1* locus in the wild type 3D1 and the insertion of a hygromycin resistance cassette *(Hyg.R)* in 3D1*Δzmr1* and in the ectopic controls. (D) Growth of 3D1Δ*zmr1* was reduced compared to growth of the ectopic controls in the presence of hygromycin (100 μg/ml). Growth is normal for all lines in the absence of hygromycin. The experiment was performed three times with similar results.

To confirm the role of transposable elements in down-regulating *Zmr1* expression, attempts were made to generate *in locus* complementation or allele replacement transformant lines of 3D1Δ*zmr1.* However, no successful transformants were obtained. Instead, we replaced the entire transposable element island (30 kb) with a hygromycin resistance cassette. Three independent knockout lines (Δ*TE*) of the transposable element insertions were obtained and analyzed for melanin accumulation *in vitro* at 7 dpi. The transposable element deletion mutants were much darker than the wild type 3D1 (Fig 6A, Additional file 8). Furthermore, expression levels of *Zmr1* in the transposable element knockouts were significantly higher than in the wild type 3D1 (Fig 6B). Overall, these results demonstrate that the transposable element island upstream of *Zmr1* in the less melanized strain negatively regulates gene expression and contributes to the variability in melanin accumulation between the two strains.

**Fig 6.**
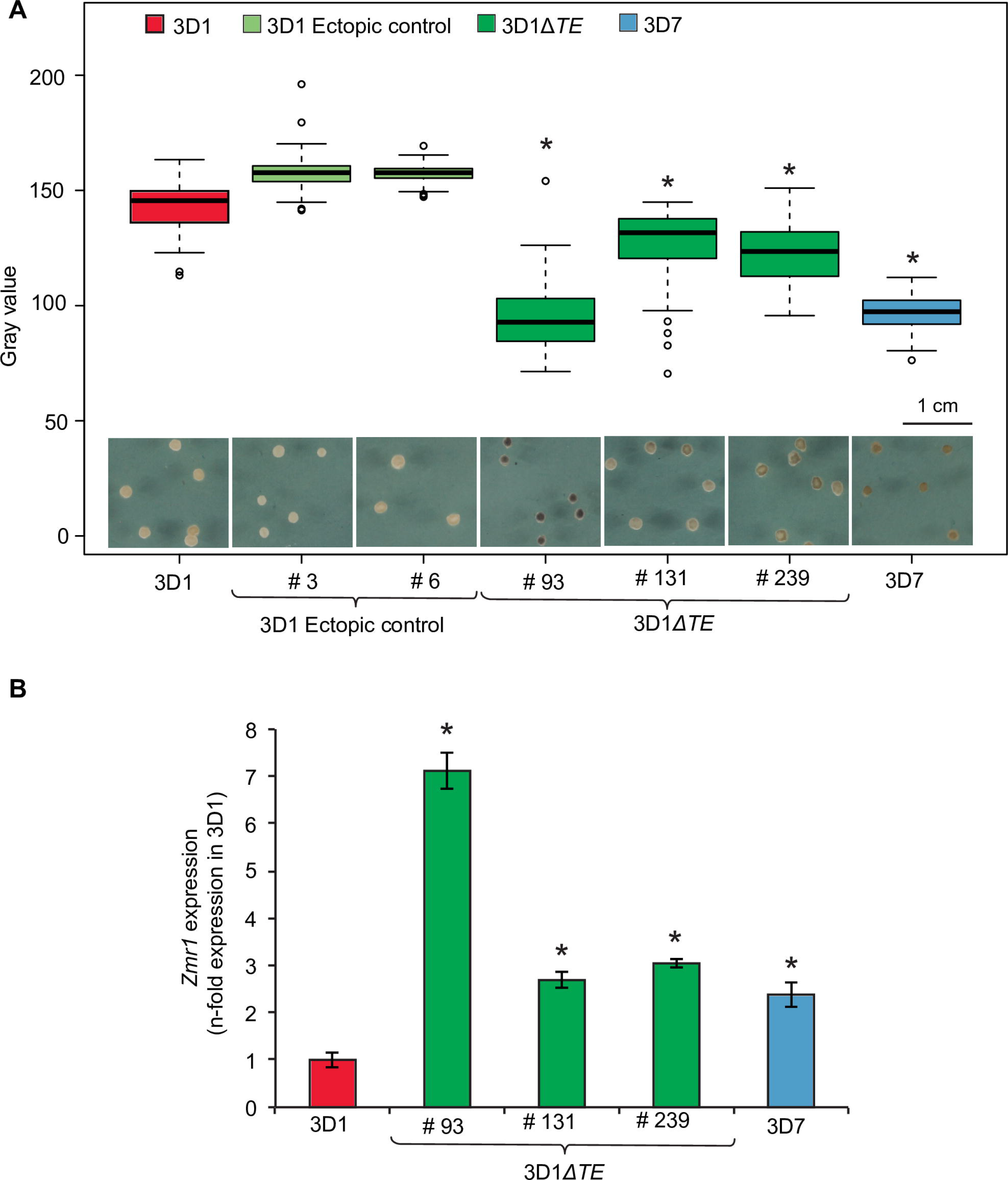
The transposable element insertion in the lighter strain down-regulates *Zmr1* expression. (A) Deletion of the transposable element island leads to significantly more melanin accumulation in 3D1 at 7 days post inoculation (dpi). Distribution of gray values for at least 35 colonies at 7 dpi for 3D1, 3D7, the transposable element deletion mutants in the 3D1 background (3D1*ΔTE* #93, #131 and #239) and the ectopic controls (#3 and #6). Asterisks indicate significant differences in gray values with respect to the wild type 3D1 (p-value ≤ 0.05, Kruskal-Wallis). (B) *Zmr1* expression levels in the transposable element knockouts (#93, #131, #239) are significantly higher than in the wild type 3D1 at 7 dpi. *Zmr1* expression values are relative to the expression of *Zmr1* in 3D1. Means and standard deviations of three technical replicates are shown.Asterisks (*) represent statistical differences with the wild type (p-value ≤ 0.05, Kruskal-Wallis test). The experiment was performed twice and we obtained similar results.

### Melanin lowers fungicide sensitivity, but has an associated trade-off

We observed that the non-melanized mutants grew faster than the corresponding wild types (Fig 7A and 7B, Additional files 9 and 10), suggesting that melanin production has a fitness cost for *Z. tritici.* In addition, we aimed to explore possible biological roles for melanin in *Z. tritici.* Virulence of a non-melanized mutant was not altered compared to the wild type strain after 21 days of infection in wheat plants (Additional files 11 and 12). Furthermore, pycnidiospores produced under controlled greenhouse conditions in the albino pycnidia of Δ*zmr1* were fully viable. Thus, we found no evidence that melanin plays a role in host colonization or pathogen reproduction. To evaluate the role of melanin in fungicide sensitivity, we grew the wild type 3D7 and the non-melanized mutant 3D7Δ*zmr1* colonies in rich media until 3D7 was melanized (5 dpi) and then we treated the colonies with the SDHI fungicide bixafen. The decrease in growth in the presence of the fungicide of the non-melanized mutant 3D7Δ*zmr1* was higher than that of the wild type 3D7, indicating that melanin lowers the sensitivity of *Z. tritici* to bixafen (Fig 7C, Additional file 13). However, the non-melanized mutant was not more sensitive to the azole fungicide propiconazole than the wild type (Additional file 13). These data demonstrate that melanin can specifically protect *Z. tritici* against SDHI fungicides, but its production has a negative effect in growth. We propose that fine-tuning of *Zmr1* expression potentially balances its beneficial functions against the growth trade-offs associated with melanin synthesis.

**Fig 7.**
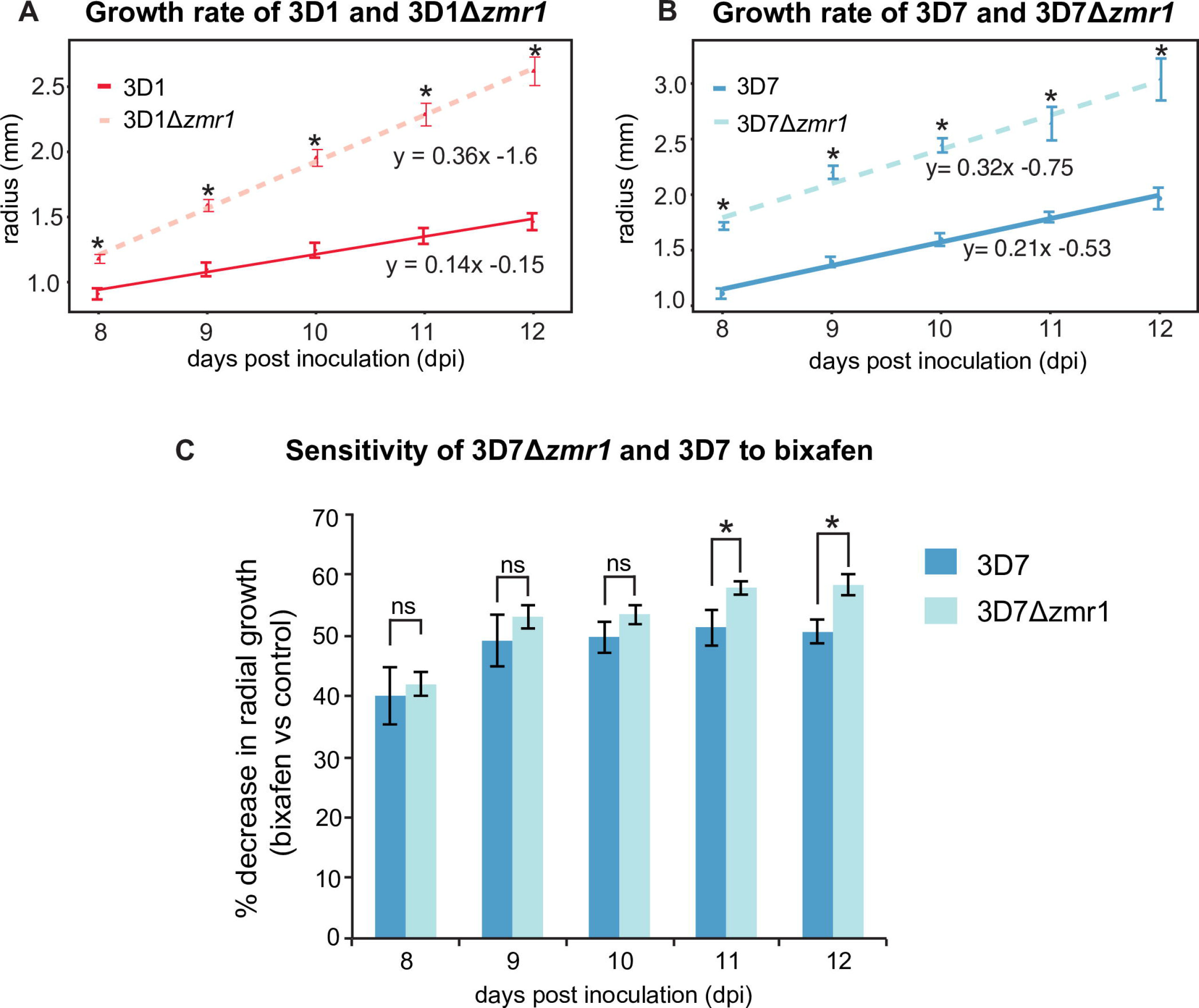
Melanin accumulation is associated with reduced growth and resistance to SDHI fungicides. (A & B) Colony radius over time of melanin deficient mutants in 3D1 (A) and 3D7 (B) backgrounds. The experiment was performed three times with similar results. (C) Melanin protects *Z. tritici* against the SDHI fungicide bixafen. Percentage decrease in growth of the wild type 3D7 and the 3D7 *Δzmr1* knockout in the presence and absence of the fungicide at each time point (8-12 dpi). Mean and standard error of differential radial size of colonies grown on three independent plates are presented. The experiment was performed twice with similar results. Asterisks (*) indicate statistical differences between wild type and knockout at each time point (p-value ≤ 0.05, Kruskal-Wallis). ns = nonsignificant.

### Transposable element insertions regulate *Zmr1* expression and melanin accumulation in *Z. tritici* populations

We hypothesized that transposable element insertions similar to those in 3D1 could contribute to differential melanization levels at the species level. We used Illumina reads from 132 *Z. tritici* strains from four different global field populations and screened for the presence of transposable elements upstream of the *Zmr1* gene. The amino acid sequence of Zmr1 was highly conserved in all the strains, with an average identity of 99%. Twelve of the strains (including 3D1) had at least one transposable element insertion within 4 kb upstream of the *Zmr1* gene. In two additional strains, short scaffold lengths prevented a full screening for the presence of transposable elements. It is likely that all of the identified insertions were the consequence of independent insertion events because they consisted of different types of transposable elements (including a retrotransposon, six DNA transposons and three unclassified transposable elements) and were located at different positions upstream of *Zmr1* (Fig 8A). We selected 11 strains with transposable element insertions and 22 without any insertion upstream of *Zmr1* to evaluate the effects of the transposable elements on the regulation of melanin production. Melanin accumulation among these strains was highly variable, with gray values ranging from 91 to 161 at 7 dpi (Fig 9, Additional files 14 and 15). Overall, insertions of transposable elements negatively affected melanin accumulation, as the strains containing insertions of transposable elements were significantly less melanized than the strains without the insertions (Fig 8B, Additional file 15). Furthermore, transposable element insertions negatively affected *Zmr1* expression levels (Fig 8C). These results further support the hypothesis that the transposable element insertion polymorphism affects *Zmr1* expression and contributes to the observed phenotypic diversity for melanin accumulation in *Z. tritici*.

**Fig 8.**
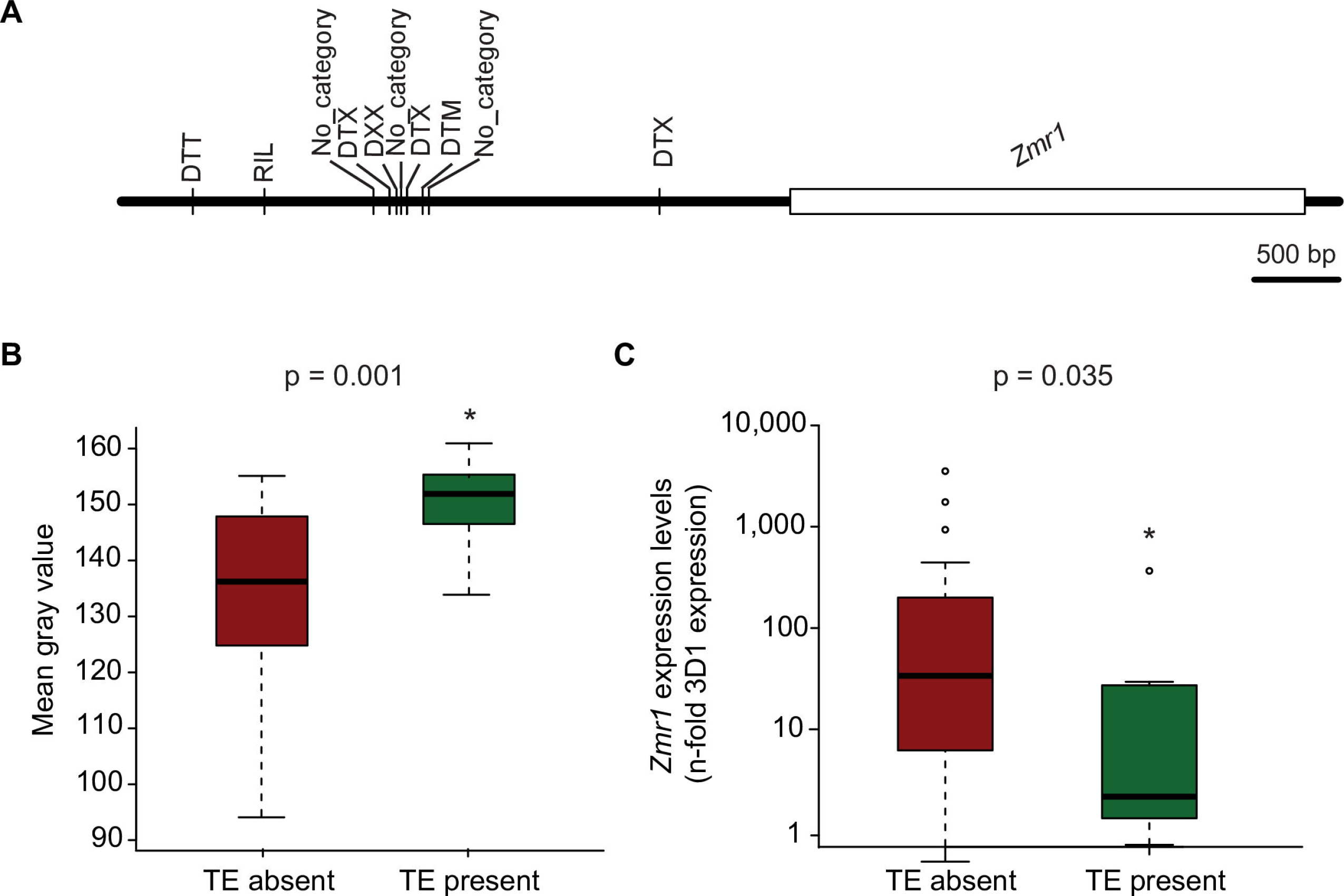
Transposable element insertions regulate *Zmrl* expression and melanin accumulation in *Z. tritici* populations. (A) Schematic representation of the location and classification of the transposable element insertions upstream of *Zmr1* in different *Z. tritici* strains from a worldwide collection. The transposable elements were classified according to the three-letter code described in Wicker *et al.*, (2007) [87]: The first letter indicates the class (R = RNA class and D = DNA class); the second letter indicates the order (I = Line, T= TIR, X = unknown); and the third letter indicates the superfamily (L = *L1*, M = *Mutator*, T = *Tc1-Mariner*, X = unknown). Transposable element insertions upstream of *Zmr1* significantly contribute to a reduction in melanin accumulation, according to Kruskal-Wallis (p-value = 0.0008, indicated with asterisks). Gray value distributions of *Z. tritici* strains with and without transposable element insertions upstream of *Zmr1.* The experiment was performed three times and we obtained similar results. (C) Transposable element insertions upstream of *Zmr1* negatively contribute to *Zmr1* expression (Kruskal-Wallis, p-value = 0.035, indicated with asterisks). Distribution of the mean expression of *Zmr1* (relative to 3D1 at 7 days post inoculation) in each *Z. tritici* strain with and without transposable element insertions upstream of *Zmr1*. The experiment was performed twice with similar results.

**Fig 9.**
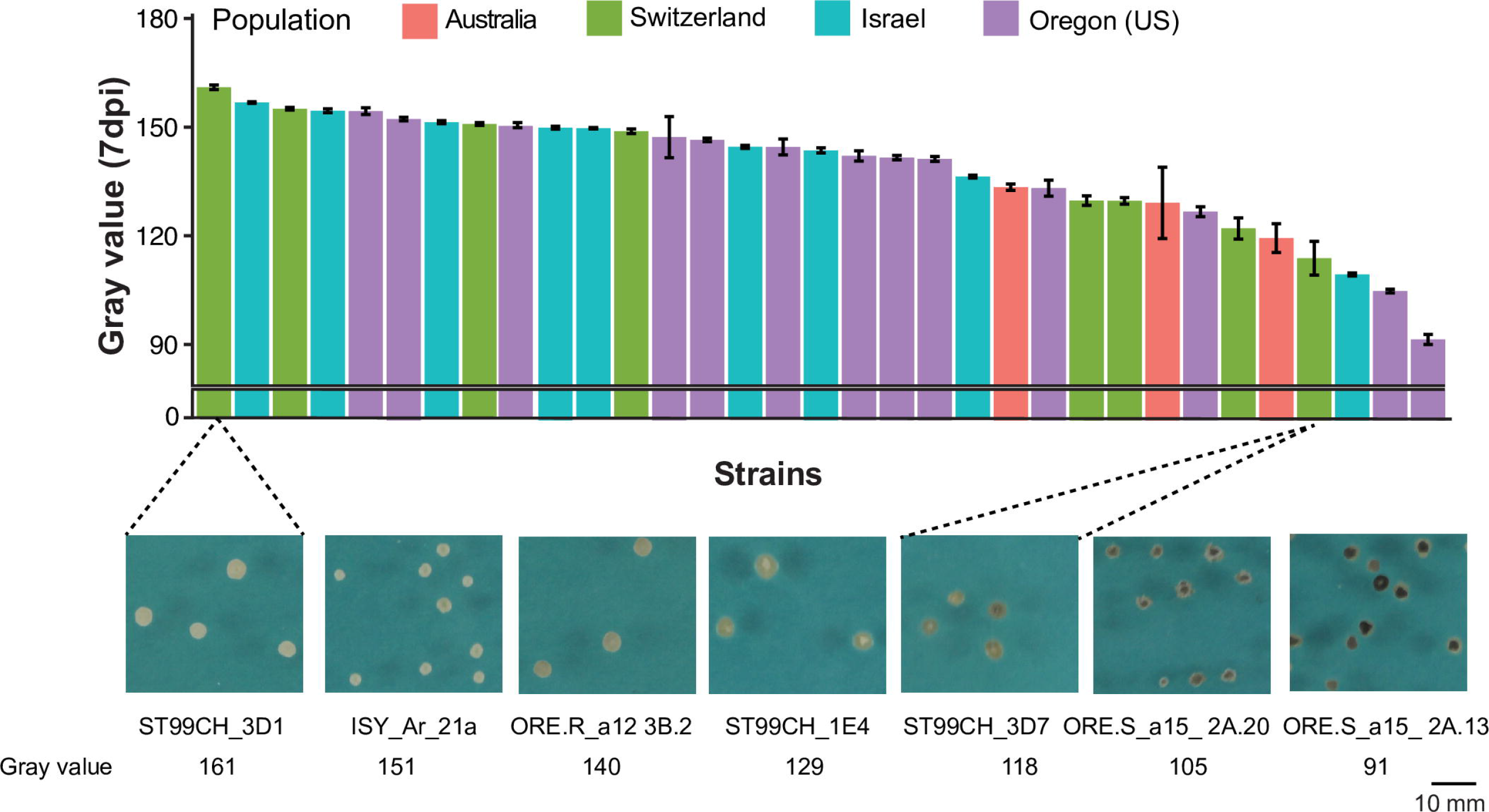
High diversity in melanin levels is exhibited among strains from four worldwide populations. Gray values of 37 different *Z. tritici* strains from four different field populations across the world. The colors of the bars indicate the population to which the strains belong. Means and standard errors of gray values were based on colonies grown for 7 days. Some examples of *Z. tritici* strains illustrating the diversity in melanin accumulation are shown in the lower panel. The experiment was performed three times with similar results.

## Discussion

Melanin is thought to play an important role in adaptation to changing environments for many fungi. Given its importance, evolution is likely to favor the emergence of genetic mechanisms that enable a finely tuned regulation of melanin accumulation that can balance fitness costs associated with melanin synthesis against the survival advantage that may be gained under hazardous conditions. Here, we demonstrated that differences in regulation of gene expression can be governed by both transposable elements and variation in promoter sequences, likely contributing to adaptive genetic variation.

### Diversity in melanin levels and its role in adaptation

Melanin is a widely distributed compound in eukaryotes that can affect fitness. The biological functions of melanin differ substantially among species [31,43–45]. In plant pathogens, such as *Pyricularia grisea, Colletotrichum lindemuthianum* and *Colletotrichum lagenarium*, melanin accumulation in the appressorium is essential for direct penetration of the host epidermis [42,46]. Because *Z. tritici* enters the host through stomata, melanin is not required to initiate infection. The lack of significant differences in virulence between isogenic melanized and non-melanized strains of *Z. tritici* suggests that melanin does not play a major role in colonization under the tested conditions. However, we cannot discount a virulence function for melanin under natural conditions, where variation in UV radiation, host genotypes and interactions with other microbes are likely to play important roles. *Z. tritici* pycnidia are highly melanized and melanin likely protects the embedded pycnidiospores. In other organisms, melanin shields against stress [43] and the degree of melanization can be correlated with the degree of resistance to stress [47]. We found that melanin can lower sensitivity to an SDHI fungicide (Additional file 13), suggesting that frequent applications of SDHI fungicides onto wheat fields may select for strains that can accumulate higher levels of melanin. The capacity of melanin to shield against toxic compounds could reflect a role for melanin in protection against antimicrobials produced under natural conditions by microbial competitors or by the host [43,48,49]. Though melanin can contribute to survival in the environment, our experiments indicated that melanin production has a fitness cost that results in reduced growth. We found that *Z. tritici* strains exhibit temporal differences in melanin accumulation. We postulate that these differences reflect selection operating to balance rates of growth with survival to environmental stress. Under this scenario, melanin accumulation illustrates how a trade-off between adaptation and growth can contribute to variation in an adaptive trait.

### Variability in melanin accumulation is caused by differential regulation of gene expression

Our approach revealed that variability in melanin accumulation is mediated by differential regulation of expression of the *Zmr1* gene. *Zmr1* encodes a transcription factor that regulates expression levels of genes in the melanin biosynthetic cluster. We characterized two regulatory layers mediating variation in *Zmr1* expression: promoter sequence modifications and an insertion of transposable elements upstream of the promoter. Twelve SNPs in the promoter of *Zmr1* underlie differential regulation of melanin accumulation in the light and dark strains. Although the individual effects of these 12 mutations have not yet been tested, we hypothesize that at least one of these promoter mutations alters the activation of *Zmr1* transcription.

An island of 13 transposable elements of approximately 30 kb is located upstream of the *Zmr1* promoter in the lighter strain and delays *Zmr1* expression. We demonstrated the contribution of the transposable elements in down-regulating melanin accumulation by removing the full transposable element island, which led to an increase in *Zmr1* expression and melanin accumulation. The transposable element-mediated down-regulation of *Zmr1* is transient, as the differences in *Zmr1* expression between the lighter and darker strain eventually reduces with age. The transposable element island hinders *Zmr1* expression either by blocking the activity of activators upstream of the transposable elements or by epigenetically silencing adjacent regions. Remarkably, we observed a silencing effect of the hygromycin resistance gene under the control of a constitutive promoter when it was located at the *Zmr1* locus, downstream of the transposable element island in the 3D1 strain. The expression of the hygromycin resistance gene was higher when it was located ectopically or at the *Zmr1* locus in the 3D7 background. These findings suggest that the transposable element insertions reduce the expression of *Zmr1* in the lighter strain through epigenetic mechanisms. Transposable elements are frequently associated with heterochromatic regions of the genome and this limits transposable element activity and transcription [40,50–54]. The spread of the heterochromatic state of the transposable elements to neighboring genes silences their expression, as shown in other organisms [15,55–58]. Frequently, under stressful conditions, some families of transposable elements are transcriptionally activated [59–61]. This suggests that transposable elements may provide a mechanism to specifically regulate expression of nearby genes under stressful conditions [21,62,63]. In *Epichloë festucae* two genes involved in the biosynthesis of alkaloids are located in a transposable element-rich region and are epigenetically silenced in axenic culture. Epigenetic silencing and de-silencing were shown to provide an important regulatory layer to specifically produce the alkaloids during host colonization [50]. In the pathogenic fungus *Leptosphaeria maculans* effector genes are located in heterochromatic regions rich in transposable elements. Insertions of transposable elements were shown to modify the epigenetic state of nearby effector genes and consequently modulate their expression patterns [64]. In maize insertion of a transposable element and the resulting spread of DNA and histone methylation marks to the cis-regulatory region of a gene reduces the accessibility for transcription factors and the RNA polymerase, thus altering expression levels upon attack by *Fusarium graminearum* [15]. We postulate that regulation of *Zmr1* by insertions of transposable elements is mediated by similar mechanisms, which involves the spreading of epigenetic marks to *Zmr1* in the lighter strain. In this way, transposable element insertions can provide a new layer of gene regulation that can optimize fitness in fluctuating environments.

### Genomic rearrangements modulate melanin levels in *Z. tritici* populations

Two antagonistic consequences of melanin accumulation, protection from stress and decrease in growth rate, suggest the need for specific regulation of melanin synthesis to enable adaptation to different environments. During host colonization, *Z. tritici* is exposed to different micro-climatic conditions and is subjected to environmental changes, depending on its spatial location during host colonization [25,65]. It is likely that this spatial and temporal environmental heterogeneity leads to diversification of melanization levels in *Z. tritici.* Fluctuations in macro-climate may also select for diversification in melanization, with episodes of severe heat, cold, drought or UV radiation likely favoring strains with higher melanization, while less melanized strains may have higher fitness during less stressful weather conditions. The significant variability in the degree of melanization exhibited among different strains of *Z. tritici* can have many underlying causes, but we hypothesize that most of these differences reflect local adaptation.

The genome of *Z. tritici* contains approximately 17% repetitive elements [39,68]. Transposable element insertions can cause adaptive variation and contribute to pathogen evolution. Transposable elements are frequently associated with stress-related genes and are considered to contribute to their diversification [22,23,63,67], but how transposable elements drive adaptation remains to be fully understood. Here we show that transposable elements contribute to phenotypic diversity by regulating gene expression. Independent insertions of transposable elements in *Z. tritici* contributed to differential regulation of *Zmr1* expression and to diversification of melanin accumulation, an important adaptive trait.

## Conclusions

We demonstrated that diversity in melanin accumulation is determined by differential regulation of gene expression instead of through mutations in coding sequences. Both single nucleotide polymorphisms in the promoter region of the *Zmr1* gene and transposable element insertions altered the accumulation of melanin. The complexity at the locus suggests that a sophisticated regulatory mechanism has evolved to balance the trade-offs between growth and melanin production. We believe that variation in transposable element insertions creates differential regulatory patterns through chromatin modification, generating new epialleles. We elucidated how transposable elements can facilitate the diversification of adaptive traits by generating regulatory variation that can fine-tune fitness-relevant gene expression.

## Methods

### Growth conditions for *Z. tritici* strains and bacterial strains

The *Z. tritici* Swiss strains ST99CH_3D1 (abbreviated as 3D1) and ST99CH_3D7 (abbreviated as 3D7) collected in 1999 [33,68] were used for genetic modifications. Wild type and genetically modified *Z. tritici* strains were grown in 50 ml of yeast sucrose broth (YSB, 1% w/v yeast extract, 1% w/v sucrose) with 50 μg/ml kanamycin sulfate in 100-ml Erlenmeyer flasks at 18°C, 120 rpm for 6 days. Blastospores from the wild type and genetically modified *Z. tritici* strains were collected after 6 days of growth in YSB. Liquid cultures were filtered through double-layered sterile cheesecloth and blastospores were collected by centrifugation (3273 g, 15 min, 4°C). The supernatant was discarded, blastospores were washed twice and re-suspended in sterile deionized water and stored on ice until use (0-1 days). The concentrations of the spore suspensions were determined using KOVA^®^ Glasstic^®^ counting chambers (Hycor Biomedical, Inc., USA). Yeast malt sucrose agar (YMS, 0.4% w/v yeast extract, 0.4% w/v malt extract, 0.4% w/v sucrose, 1.5% w/v agar) and potato dextrose agar (PDA) were used for growing Z *tritici* strains on Petri plates.

*E. coli* strains NEB^®^ 5-alpha (New England Biolabs) or HST08 (Takara Bio, USA) were used for molecular cloning. *E. coli* strains were grown on DYT media (1.6% w/v tryptone, 1% w/v yeast extract, 0.5% NaCl) amended with kanamycin sulfate (50 μg/mL) at 37°C. *Agrobacterium tumefaciens* strain AGL1 was used for *A, tumefaciens-m*ediated transformation of *Z. tritici. A. tumefaciens* was grown in DYT media containing kanamycin sulfate (50 μg/ml), carbenicillin (100 μg/ml) and rifampicin (50 μg/ml) at 28°C, unless stated otherwise.

### QTL mapping

Phenotypic data (gray values of the mapping population at 8 dpi) and RADseq data from the progeny of the cross between 3D1 and 3D7 described earlier [33] was used for QTL mapping, using the same protocol described in Meile *et al.,* 2018 [69]. QTL re-mapping of chromosome 11 alone was performed in R/qtl version v1.40-8 [70] by simple interval mapping (SIM) analysis as described previously [33].

### Melanization analysis

The degree of melanization in each *Z. tritici* strain was estimated by plating approximately 100 blastospores on YMS plates. Plates were then randomized and incubated in the dark at 22°C and 70% humidity. Digital images of the plates were taken through the Petri plate lid at different time points, using standardized settings [33]. Gray value, a proxy for degree of melanization, was estimated for each colony using ImageJ [71]. The gray scale ranges from 0 to 255, where 0 represents the darkest shade of black and 255 represents the lightest shade of white. Gray value of colonies grown on at least three independent Petri plates was measured.

### Measurements of growth rate and fungicide sensitivity assays

Because the 3D7Δ*zmr1* mutant grew as hyphae instead of as blastospores in YMS (Additional file 16), it was not possible to make a proper evaluation of its growth rate on YMS. Thus, we performed these experiments on PDA, on which both knockouts grew with a morphology that was similar to the wild type strains (Additional file 16). Colony size was evaluated as previously described at 7-12 dpi [30]. The growth curve for wild type strains and knockouts was obtained by plotting radial growth (mm) over time and fitted to a linear model (Pearson’s correlation coefficient value (r^2^ value > 0.9). Growth rate (mm/day) was estimated by calculating the slope of the growth curve. Analysis of covariance (ANCOVA) was performed to determine if there were significant differences in growth rate (p-value ≤ 0.05). Significant differences in colony size at each time point (Kruskal-Wallis, ≤ 0.05) were evaluated between Δ*zmr1* and the wild type. The experiment was performed three times. To perform fungicide sensitivity assays comparing the wild type 3D7 and non-melanized 3D7Δ*zmr1* line, a 100-blastospore suspension was plated on sterile Whatman filter paper, grade 1 (Huber lab) placed on PDA plates. Three plates per strain and condition were incubated in the dark at 22°C with 70% humidity. After 5 days the plates were photographed and the filter papers were transferred to PDA plates supplemented with fungicides (0.75 ppm of bixafen or 0.75 ppm of propiconazole, Syngenta, Basel, Switzerland) or control PDA plates without any fungicides. The strains were grown under the same conditions as before and digital images were captured every 24h for 8-12 days. The radial growth rates were calculated as described earlier using ImageJ [41]. The percentage decrease in colony radius in the presence of each fungicide was calculated at each time point. The experiment was performed twice.

### Generation of *Z. tritici* transformants

All the amplifications were performed using Phusion high-fidelity DNA polymerase from NEB (Ipswich, MA, USA). *Zmr1* disruptant mutants in both 3D1 and 3D7 backgrounds were generated by inserting a hygromycin resistance cassette into the *Zmr1* gene 13 base pairs (bp) after the start codon using homologous recombination (Additional file 17). Up-flanking and down-flanking regions (approximately 1000 bp) of the site of integration were PCR-amplified from either 3D1 or 3D7 genomic DNA. A hygromycin resistance cassette with the desired overlap for In-Fusion cloning was amplified from the plasmid pES6 (obtained from Eva Stukenbrock, Kiel University). The flanking regions and the hygromycin resistance cassette were fused to the binary vector backbone of pES1 (obtained from Eva Stukenbrock, Kiel University) in their respective order (Additional file 17) by a single step In-Fusion reaction (TakaraBio, Mountain View, CA, USA) following the manufacturer’s instructions and then cloned in *E. coli.*

Constructs to generate knockouts of the transposable elements in the 3D1 background were obtained in a similar way, except that these mutants were generated by replacing the transposable elements by the hygromycin resistance cassette (Additional file 17).

For generating *in locus* allele swaps and complementation lines, the full-length *Zmr1* gene along with 1863 and 1862 bp upstream of the start codon in 3D7 and 3D1, respectively and 539 bp downstream of the stop codons, were amplified and fused to a geneticin resistance cassette amplified from the pCGEN vector [72] and the vector backbone of pES1 as described earlier (Additional files 17 and 18). This intermediate construct was used to amplify the full *Zmr1* gene fused to the geneticin resistance cassette. Additionally, approximately 1 kb upstream and downstream of the insertion site in 3D7 were amplified and the three amplicons were fused to the binary vector backbone of pES1 as described earlier (Additional files 17 and 18).

Mutation-free plasmids were transformed into the *A. tumefaciens* strain AGL1 [73] by electroporation and screened on DYT medium supplemented with 50 μg/ml rifampicin, 50 μg/ml carbenicillin and 40 μg/ml kanamycin at 28°C. *A. tumefaciens-mediated* transformation of *Z. tritici* was performed as previously described [69,74,75]. Selection of transformants was performed on YMS plates containing 200 μg/ml cefotaxime and the corresponding antibiotic, either hygromycin at 100 μg/ml (Neofroxx, Germany) or geneticin at 150 μg/ml (Thermo Fisher Scientific) at 18°C for 8-12 days. Individual colonies were then streaked onto YMS plates containing the corresponding antibiotic and grown at 18°C for one week. After one round of selection single colonies were transferred to YMS plates without a selection agent and transformants were screened for the correct inserts by colony PCR using KAPA3G Plant DNA polymerase (Kapa Biosystem, Massachusetts, USA) and specific primers (Additional file 18). These amplicons were further sequenced (Microsynth AG, Balgach, Switzerland) to confirm the correct integration. The copy number of the transformants was determined by performing quantitative PCR (qPCR) on DNA isolated from transformed Z *tritici* strains using Qiagen plant DNeasy kit (Qiagen) and specific primers for the antibiotic resistance marker and for the housekeeping genes *TFIIIC1* or *18s rRNA* (Additional file 18), as previously described [69]. DNA from wild type *Z. tritici* strains without the transgene, DNA from *Z. tritici* strains harboring a single transgene and negative water controls were included in all analyses.

### Hygromycin resistance assay

To test the sensitivity to hygromycin of 3D1Δ*zmr1* and 3D7Δ*zmr1*, their respective wild types and the ectopic controls, 5 μl of 10^4^ spores/ml of 6-day old blastospores were drop inoculated on YMS media supplemented with hygromycin at 100 μg/ml (Neofroxx, Germany). YMS media without hygromycin was used as a control. Images were taken at 8 dpi.

### Comparative transcriptomic analysis

RNA sequencing (RNA-seq) analysis was performed to identify differentially expressed genes in wild type and melanin deficient Δ*zmr1* mutants. Roughly 100 blastospores of *Z. tritici* strains 3D1, 3D1 Δzmr1 #6, 3D7, and 3D7Δzmr1 #48 were placed onto PDA plates and incubated at 22°C in the dark with 70% humidity. After 7 days individual colonies were picked carefully from the plates using sterile forceps, collected and frozen in liquid nitrogen. The colonies were then homogenized using a Bead Ruptor with a cooling unit (Omni International) and zirconium oxide beads (1.4 mm). RNA was extracted using the GENEzol reagent (Geneaid Biotech) following the manufacturer’s recommendations. On column DNAase treatment was performed using RNeasy mini kit (Qiagen) following the manufacturer’s instructions.

RNA-seq was performed on an Illumina HiSeq 2500 using paired end reads at 2 × 101 bp as previously described [1]. Raw RNA-seq reads were trimmed using Trimmomatic v. 0.33 [76]. Trimmed reads were aligned to the *Z. tritici* parental genome 3D7 or 3D1 and transcriptome using TopHat v 2.0.13 [77]. Gene counts were calculated using HTSeq v0.6.1 [78] and differential gene expression analysis was performed using the R package EdgeR version 3.2.3 [79]. Relative RNA levels in the RNA-seq experiment were calculated by TMM (trimmed mean of M-values) normalization [80]. Mean TMM normalized log_2_ CPM (counts per million mapped reads) were calculated for all the annotated genes. To identify differentially expressed genes between a wild type strain and melanin-deficient Δ*zmr1* knockouts, Benjamin-Hochberg false discovery rates (FDR) and an FDR adjusted p-value were calculated. The RNAseq was deposited in SRA database with the accession number SRP143580 (https://www.ncbi.nlm.nih.gov/sra/SRP143580).

### Quantitative reverse transcription PCRs (qRT-PCRs)

Expression levels of *Zmr1* in different *Z. tritici* strains and genetically modified strains were quantified using qRT-PCR. RNA was extracted from *Z. tritici* strains grown *in vitro* and harvested at 7 dpi or 9 dpi depending on the experiment, as described earlier for the RNA-seq analysis. cDNA was synthesized from 500 ng of RNA using oligo(dT)_18_ primers and Revert Aid RT Reverse Transcription kit (Thermo Scientific) following the supplier’s instructions. qRT-PCR analysis was performed using a 10-μl reaction mix with 1 μl of cDNA. A negative control with RNA alone and water was also included. Specific primers spanning introns were designed for the targets *Zmr1* and 18S ribosomal RNA to avoid the risk of genomic DNA contamination (Additional file 18). Crossing point (Cp) values were calculated using absolute quantification and the second derivative method provided by LightCycler 480 software version 1.5 (Roche Diagnostics Corp., Indianapolis, IN, USA). “Advanced Relative Quantification” method was used to analyze the fold change in expression of *Zmr1* when compared to the wild type strains. “Advanced Relative Quantification” method was also used for estimating the fold change in expression of *Zmr1* in different strains of *Z. tritici* compared to 3D1.

### *In planta* virulence assay

The *Z. tritici* wild type 3D7 strain and three independent 3D7Δ*zmr1* mutants lacking melanin were compared for their ability to infect the winter wheat (*Triticum aestivum*) variety Drifter. Two wheat seeds were sown in peat soil (Jiffy GO PP7, Tref, Netherlands) in 7 × 7 × 9 cm plastic pots and grown in a greenhouse at 18°C day and 15°C night, with a 16 h light cycle and 70% relative humidity. Plants were fertilized 10 days after sowing with 10 ml 0.1% Wuxal Universaldünger (Maag AG, Switzerland) per pot. Twelve seventeen day-old seedlings were spray-inoculated with 15 ml of a blastospore suspension (10^6^ spores/ml) containing 0.1% (v/v) of Tween 20 (Sigma Aldrich). Pots were placed under 100% humidity for 3 days by covering them with a plastic bag. The second leaf of each plant was collected at 21 dpi and pycnidia density (pycnidia/cm^2^ leaf) and percentage of leaf area covered by lesions (PLACL) was analyzed using automated image analysis which was manually verified [37].

### Annotation of transposable elements in *Z. tritici* strains and sequence alignment

For *Z. tritici* strains 3D1, 3D7, 1E4 and 1A5 full genome annotations were already available [39,81]. We annotated and masked repetitive elements for the remaining 128 *Z. tritici* strains using RepeatModeler version 1.0.8 as described earlier [39,69]. We masked the genomes using RepeatMasker version 4.0.5 with the library previously obtained for *Z. tritici* strain IPO323 [38] according to the transposable element nomenclature defined by Wicker *et al.*, (2007) [82]. Multiple sequence alignment of *Zmr1* in the *Z. tritici* strains was performed using AliView version 1.22 [83]. Amino acid sequence identity of Zmr1 in the *Z. tritici* strains was calculated using the Sequence Identities and Similarities (SIAS) [84].

### Statistical analysis

Data analyses and plotting were performed using R version 3.3.1 and RStudio version 1.0.143 [85,86] and microsoft excel. The non-parametric Kruskal-Wallis test was used to compare gray values between different strains/groups. Tukey’s HSD test was used to estimate significant differences in pycnidial density between different *Z. tritici* strains for the *in planta* virulence assay.

## List of abbrevations

QTL: quantitative trait locus
Zmr1: Zymoseptoria melanin regulation 1
SNP: Single nucleotide polymorphism
Pks1: Polyketide synthase 1

## Declarations

- Ethics approval and consent to participate: Not applicable
- Consent for publication: Not applicable
- Availability of data and material: The datasets generated and analysed during the current study are available in the SRA database (accession number: SRP143580), https://www.ncbi.nlm.nih.gov/sra/SRP143580

## Competing interest

The authors declare that they have no competing interests.

## Funding

The research was supported by the Swiss National Science Foundation (Grant 31003A_155955, http://www.snf.ch/en/Pages/default.aspx.) and by the ETH Zurich Research Commission (Grant 12-03, https://www.ethz.ch/en/the-ethzurich/organisation/boards-university-groups-commissions/research-commission.html) to BAM. PK was supported by the Swiss State Secretariat under the Education Research and Innovation, through the Federal Commission for Scholarships for Foreign Students. CP was supported by an INRA Young Scientist grant. DC is supported by the Swiss National Science Foundation (grant 31003A_173265, http://www.snf.ch/en/Pages/default.aspx). The funders had no role in the design of the study, data collection, analysis and interpretation of data and in writing the manuscript.

## Authors’ contributions

PK contributed to the conception and design of the analysis, acquisition of data, analysis of the data and writing the original draft of the manuscript. LM contributed to the design of the analysis and acquisition of data; contributed to the edition and revision of the manuscript. XM analyzed the RNAseq data and contributed to the edition of the manuscript. CP contributed to the analysis of the transposable elements in the populations. FEH contributed to the bioinformatics analysis of transposable elements variation in the populations and edited the manuscript. DC contributed to the analysis of the transposable elements in the populations and edited and reviewed the manuscript. BAM acquired funding and reviewed the manuscript. ASV contributed to the conception and design of the analysis, supervision and analysis of the data and writing of manuscript.

## Acknowledgements

RNA sequencing was performed at the quantitative genomics facility of the D-BSSE, ETH Zurich. qPCR was performed at the Genetic Diversity Centre (GDC), ETH Zurich. Technical assistance was provided by Lotta Köppel. Eva H. Stukenbrock and Jason J. Rudd provided us with vectors.

## Addditional files

**Additional file 1** (Additional file 1.pdf) **3D7 accumulates more melanin than 3D1**. Means and standard errors of gray values (0 = black, 255 = white) of at least 60 colonies of *Z. tritici* strains 3D1 and 3D7 at 7-12 days post inoculation (dpi). The experiment was performed three times with similar results. Asterisks indicate significant differences according to Kruskal-Wallis test (p-values ≤ 0.05).

**Additional file 2** (Additional file 2.pdf) **Comparison of the QTLs obtained using the genome of the reference strain IPO323 and of the parental strain 3D7**.

**Additional file 3** (Additional file 3.pdf) **Genes in the 95% confidence interval of the QTL obtained using the genome of the parental strain 3D7**. Genes in the *Pks1* gene cluster are indicated in bold. The total number of synonymous (Syn) and non-synonymous (Non-Syn) SNPs between both parental strains is indicated. Insertions upstream of the coding sequence are also indicated.

**Additional file 4** (Additional file 4.pdf) ***Zmr1* expression levels are lower in 3D1 than in 3D7 at 7 days post inoculation**. Mean and standard errors (se) of the relative quantification (RQ, fold change in expression level of *Zmr1* with respect to the *Zmr1* expression levels of the strain 3D1 at 7 days post inoculation, dpi) of the expression of *Zmr1* in 3D1 and 3D7 at 7 and 9 dpi. The experiment was performed three times with similar results.

**Additional file 5** (Additional file 5.tiff) **Zmr1 regulates melanin biosynthesis in *Z. tritici***

(A) Three independent *Zmr1* disruptant mutants in the 3D1 (#46, #48 and #2.1) and in the 3D7 (#3, #6 and #100) backgrounds lack melanin. Pictures of 10 day old wild type 3D7 and 3D7Δ*zmr1* colonies. (B) Melanized and albino pycnidia of 3D7 and 3D7Δ*zmr1*, respectively, on wheat leaves of the cultivar Drifter, 21 dpi (C) Expression values of the genes in the DHN melanin biosynthesis pathway (Pks1 = polyketide synthase 1; Thr1 = 1,3,8-trihydroxynaphthalene reductase; Zmr1 = *Zymoseptoria* melanin regulation 1; Ayg1 = Homolog of *Aspergillus* yellowish green) for the wild type and *μzmr1* in the 3D1 and 3D7 backgrounds, respectively. Mean of TMM (trimmed mean of M-values) normalized log_2_ CPM (counts per million mapped reads) values of three independent replicates with their standard deviation are plotted. Asterisks indicate statistical differences between the wild type and the mutant (p-value ≤ 0.05, FDR ≤ 0.05).

**Additional file 6** (Additional file 6.pdf) **Reduced expression of genes in the *Pks1* cluster in Δ*zmr1* mutants**. Mean and standard error (se) of CPM (counts per million mapped reads) values of genes significantly down regulated (false discovery rates, FDR ≤ 0.05) in both *Δzmr1* mutants compared to the wild type strains 3D1 and 3D7. Means and standard errors of three independent replicates are indicated. Genes previously shown to be involved in melanin biosynthesis are shown in bold.

**Additional file7** (Additional file 7.pdf) **Sequence variation in *Zmr1* promoter contributes to differences in melanin accumulation**. Means and standard errors of gray values (0 = black, 255 = white) of 3D1, 3D7, 3D7*Δzmr1*, 3D7Δ*zmr1* + Zmr13D1 and 3D7Δ*zmr1*+Zmr1_3D7_, 7 days post inoculation (dpi) based on at least 20 colonies. Asterisks (*) and pluses (+) indicate significant differences in mean gray values of each strain with respect to the mean gray value of 3D7 and 3D7Δ*zmr1*, respectively (Kruskal-Wallis, p-values ≤ 0.05). The experiment was performed twice with similar results. NA = not applicable.

**Additional file 8** (Additional file 8.pdf) **The transposable element insertion upstream of *Zmr1* in 3D1 down-regulates *Zmr1* expression**. Mean gray values (0 = black, 255 = white) based on at least 35 colonies of the wild types 3D1 and 3D7, three independent TE deletion mutants in the 3D1 background (3D1 *ΔTE* #93, #131, #239) and the two ectopic controls (3D1+ Hyg #3 and #6), 7 days post inoculation. sAsterisks (*) indicate that the strains are significantly darker than the wild type 3D1 (Kruskal-Wallis, p-value ≤ 0.05).

**Additional file 9** (Additional file 9.pdf) **Non-melanized mutants grow faster than the wild types**. Radial growth rates (slope of the curves) of wild type strains and melanin deficient mutants (3D1*Δzmr1* and 3D7*Δzmr1*) were obtained by plotting radial size (mm) of the colonies over time. The fit of the radial growth curve estimated using a linear model was evaluated using Pearson’s correlation coefficient (r^2^ value). Asterisks (*) indicate significant differences in growth rate (slope) according to ANCOVA analysis (p-values ≤ 0.05). The experiment was performed three times with similar results.

**Additional file 10** (Additional file 10.pdf) **Colonies of non-melanized mutants are bigger than those of the corresponding wild types**. Means and standard errors of the radial size (mm) based on at least 20 colonies at different days post inoculation (dpi). The experiment was performed three times with similar results.

**Additional file 11**. (Additional file 11 tiff) **Melanin is not essential for virulence**

All three 3D7*Δzmr1* mutants were equally virulent on the winter wheat variety Drifter, compared to the wild type 3D7 as indicated by pycnidia/cm^2^ leaf 21 days post infection (Turkey’s HSD test, p-value ≤ 0.05). Mean and standard error of 12 independent leaves are shown. The experiment was performed three times for line #6 with similar results.

**Additional file 12** (Additional file 12.pdf) **Melanin deficient mutants are not impaired in virulence**. Mean of percentage of leaf area covered by lesions (PLACL) and pycnidia/cm^2^ lesion caused by the wildtype 3D7 and 3D7*Δzmr1* lines on the wheat cultivar Drifter, at 22 days post inoculation. Mean and standard error of 12 leaves are shown. The experiment was performed twice with the line 6 and 3D7 obtaining similar results. No statistical differences in PLACL and pycnidia/cm^2^ leaf were detected between 3D7 and 3D7*Δzmr1* according to Tukey’s HSD (honest significant difference) test (p-values ≤ 0.05).

**Additional file 13** (Additional file 13.pdf) **Melanin protects *Z. tritici* against SDHI fungicides**. Reduction in growth by fungicides (% growth decrease) of the wild type 3D7 and 3D7*Δzmr1*. Strains were grown on Whatman filter paper placed on PDA for 5 days and later transferred to PDA plates supplemented with the fungicide bixafen or propiconazole. Asterisks (*) indicate that the percentage in decrease in growth of the mutant strain, in the presence of the fungicide, is significantly different from the wild type 3D7 (Kruskal-Wallis, p-value ≤ 0.05). Mean and standard error of differential radial size of at least 25 colonies grown on three independent plates are presented. The experiment was performed twice with similar results.

**Additional file 14** (Additional file 14.tiff) **No significant difference in the distribution of mean gray values of different *Z. tritici* strains belonging to four different populations**

Distribution of mean gray values of different *Z. tritici* strains from four different populations across the world (3 strains from Australia, 8 from Switzerland, 9 from Israel and 13 from the US). The colors of the violin plot indicate different populations. Black dots represent individual data points, which correspond to the mean gray values of each *Z. tritici* strain at 7 days post inoculation. At least 100 colonies grown on five different plates were evaluated. The experiment was performed three times with similar results. No statistically significant differences were observed between the populations (Kruskal-Wallis test, p-value ≤ 0.05).

**Additional file 15**(Additional file 15.pdf) **Melanization and *Zmr1* expression levels in *Z. tritici* strains from around the world**. Means and standard errors of gray values (0 = black, 255 = white) and of *Zmr1* expression at 7 days post inoculation. Gray values were measured in three independent experiments, and similar results were obtained. Expression analysis was performed twice and provided similar results. AUS: Australia, CH: Switzerland, ISY: Israel and OR: USA (Oregon), TE = transposable element, P = Present, A = Absent.

**Additional file 16** (Additional file 16.tiff) **Altered growth morphology of 3D7Δ *zmr1* mutants on YMS but not in PDA**

Morphology of 3D1, 3D7 and the mutants in *Zmr1* grown on YMS and PDA at 7 days post inoculation.

**Additional file 17** (Additional file 17.pdf) **Generation of mutants** (A) Schematic diagram showing the location of primers used for generating Z*mr1* disruptant mutants (U2 + U3, D1 + D2) and the primers used for screening the transformants (U1 + Hyg UF, Hyg DF + D3). (B) Schematic diagram showing the location of primers used for generating the transposable element (TE) deletion mutants in 3D1 background (*ΔTE*; TE_U2 + TE_U3, TE_D1 + TE_D2) and the primers used for screening the transformants (TE_U1 + Hyg UF, Hyg DF + TE_D3). (C) Diagrams showing the location of primers used for generating the transformants expressing *Zmr1* gene *in locus* in the 3D7Δ*zmr1* background. Amplification of *Zmr1* gene from genomic DNA of 3D1 and 3D7 strains was performed using *Zmr1* F and *Zmr1* R primers; the geneticin resistance cassette (Gen) was amplified from pCGEN with Gen F and Gen R. Both amplicons were fused to the pES1 backbone to generate an intermediate construct, which was used to amplify *Zmr1* and the geneticin resistance cassette using *Zmr1 in locus* F + *Zmr1 in locus* R. Up-flanking and down-flanking regions of the insertion site were amplified from 3D7 genomic DNA using the primers: UF_F + UF_R and DF_F + DF_R. Primers used for screening the transformants (P1 + P2, P3 + P4) are also shown. LB and RB indicates the left and right border of the binary vector pES1. Hyg.R = hygromycin resistance cassette.

**Additional file 18** (Additional file 18.pdf) **List of primers used in the study, their sequence and their purpose**

